# Regional and hemispheric susceptibility of the temporal lobe to FTLD-TDP type C pathology

**DOI:** 10.1101/847582

**Authors:** V. Borghesani, G. Battistella, M.L. Mandelli, A. Welch, E. Weis, K. Younes, J. Neuhaus, L.T. Grinberg, W. M. Seeley, S. Spina, B. Miller, Z. Miller, M.L. Gorno-Tempini

## Abstract

Post-mortem studies show that focal anterior temporal lobe (ATL) neurodegeneration is most often caused by frontotemporal lobar degeneration TDP-43 type C pathology. Clinically, these patients are described with different terms, such as semantic variant primary progressive aphasia (svPPA), semantic dementia (SD), or right temporal variant frontotemporal dementia (FTD) depending on whether the predominant symptoms affect language, semantic knowledge for object or people, or socio-emotional behaviors. ATL atrophy presents with various degrees of lateralization, with right-sided cases considered rarer even though estimation of their prevalence is hampered by the paucity of studies on well-characterized, pathology-proven cohorts. Moreover, it is not clear whether left and right variants show a similar distribution of atrophy within the ATL cross-sectionally and longitudinally.

Here we study the largest cohort to-date of pathology-proven TDP-43-C cases diagnosed during life as svPPA, SD or right temporal variant FTD. We analyzed clinical, cognitive, and neuroimaging data from 30 cases, a subset of which was followed longitudinally. Guided by recent structural and functional parcellation studies, we constructed four bilateral ATL regions of interest (ROIs). The computation of an atrophy lateralization index allowed the comparison of atrophy patterns between the two hemispheres. This led to an automatic, imaging-based classification of the cases as left-predominant or right-predominant. We then compared the two groups in terms of regional atrophy patterns within the ATL ROIs (cross-sectionally) and atrophy progression (longitudinally).

Results showed that 40% of pathology proven cases of TDP-43-C diagnosed with a temporal variant presented with right-lateralized atrophy. Moreover, the findings of our ATL ROI analysis indicated that, irrespective of atrophy lateralization, atrophy distribution within both ATLs follows a medial-to-lateral gradient. Finally, in both left and right cases, atrophy appeared to progress to the contralateral ATL, and from the anterior temporal pole to posterior temporal and orbitofrontal regions.

Taken together, our findings indicate that incipient right predominant ATL atrophy is common in TDP-43-C pathology, and that distribution of damage within the ATLs appears to be the same in left- and right- sided variants. Thus, regardless of differences in clinical phenotype and atrophy lateralization, both temporal variants of FTD should be viewed as a spectrum presentation of the same disease.

**Highlights:** ⍰ Anterior temporal lobe (ATL) degeneration is most often caused by FTLD-TDP type C pathology

⍰ Cases can present with predominantly left (60%) or right (40%) ATL atrophy

⍰ Within ATLs, medial regions are more vulnerable than lateral ones

⍰ The observed spectrum of clinical phenotypes is driven by atrophy lateralization

⍰ Left and right temporal variants of FTD should be considered the same disease

## 1. Introduction

Frontotemporal dementia (FTD) is an umbrella term covering several clinical phenotypes associated with a progressive decline of executive functions, motor abilities, behavior, and/or language (Neary, Snowden, Gustafson, Passant, Stuss, Black, Freedman, Kertesz, Robert, Albert, Boone, et al., 1998). These clinical syndromes arise from neurodegeneration of cortical and subcortical structures within frontal and/or temporal lobes (frontotemporal lobar degeneration or FTLD) and are associated with diverse molecular pathologies (Miller et al., 1991). Converging lines of research associate the early-stages of neurodegenerative diseases to relatively focal atrophy affecting specifically susceptible cell assemblies, later spreading throughout large-scale networks (Brown et al., 2019; Raj, Kuceyeski, & Weiner, 2012; Seeley, Crawford, Zhou, Miller, & Greicius, 2009; Zeighami et al., 2015). Thus, careful clinicopathological investigations of well-defined groups of patients with focal neurodegeneration can deepen our understanding of regional vulnerability to proteinopathies (Soto & Pritzkow, 2018; Walsh & Selkoe, 2016), and have important implications for both clinical practice and cognitive neuroscience (Elahi & Miller, 2017).

Within the FTD family, in-vivo clinical studies have isolated a spectrum of syndromes characterized by selective anterior temporal lobe (ATL) atrophy, yet remarkable heterogeneity of linguistic and/or behavioral difficulties. Post-mortem, this focal anterior temporal degeneration is found to be reliably associated with abnormal depositions of the transactive response DNA-binding protein Mr43kD43 (TDP-43) type C (Rohrer, Gennatas, & Trojanowski, 2010; Spinelli et al., 2017). Ante-mortem, patients might be diagnosed with semantic dementia (SD (Julie Snowden, Goulding, & Neary, 1989)), semantic variant PPA (svPPA, (Gorno-tempini et al., 2011)), right temporal variant FTD (Thompson, Patterson, & Hodges, 2003) or behavioral variant FTD (bvFTD, (Rascovsky et al., 2011)) according to the clinical criteria adopted and the neuropsychological tests performed. Clinical symptoms in these patients might affect language, behavior, and/or emotional processing: e.g., anomia and word comprehension deficits, inability to recognize objects and famous faces, disinhibition, and facial expression recognition difficulties. While most patients present with bilateral atrophy and overlapping language and behavioral symptoms, historically two main clinical profiles have been described in relation to atrophy lateralization. Cases with predominantly left atrophy have been associated with greater naming, word comprehension, reading and object semantic deficits (Lambon Ralph, McClelland, Patterson, Galton, & Hodges, 2001; Marsel Mesulam et al., 2013). These patients almost invariably meet current consensus criteria for svPPA (Gorno-tempini et al., 2011). On the other hand, the clinical phenotype of right ATL atrophy is less well defined and has been associated with various degrees of socio-emotional (Edwards-Lee et al., 1997; Perry et al., 2001; Rosen et al., 2002; Rankin et al., 2006) and nonverbal semantic impairments (e.g., person-specific knowledge; Chan et al., 2009; Snowden et al., 2017; Woollams & Patterson, 2017). These patients might not clearly meet criteria for overall PPA (McCarthy & Warrington, 2016), and their nonverbal semantic loss can often be captured only with specialized neuropsychological tests (e.g., identification of famous faces). Overall, fewer right-predominant cases have been described (Hodges et al., 2010), but attempts to estimate the prevalence of left vs. right ATL involvement in FTLD-TDP-43-C have been hampered by the paucity of studies on pathology-proven datasets. Furthermore, misclassifications are likely when diagnoses are based on clinical criteria that have not been tailored to include nonverbal semantic deficits. For instance, it is currently hard to distinguish, from history and neuropsychological testing, the right temporal variant of svPPA/SD patients from bvFTD ones, as they might both present with a range of emotional and behavioral changes. Early, reliable differential diagnosis between svPPA and bvFTD is critical because the latter is not associated with a predominance of TDP-43-C pathology but rather with a variety of FTLD subtypes (Perry et al., 2017). To date, our understanding of the right temporal variants has relied on data stemming from single cases (e.g., (Barbarotto, Capitani, Spinnler, 1995; Gainotti, Barbier, & Marra, 2003; Joubert et al., 2004; Mendez & Ghajarnia, 2001)), or small samples without pathological diagnosis (e.g., (Miller, Chang, Mena, Boone, & Lesser, 1993; Edwards-Lee et al., 1997, Seeley et al., 2005; Binney et al., 2016; Brambati et al., 2009; Chen et al., 2018; Hodges et al., 2010; Kumfor et al., 2016; Mion et al., 2010; Snowden, Thompson, & Neary, 2012; Snowden et al., 2017; Thompson et al., 2003; Woollams & Patterson, 2017)). Here, we describe and compare the clinical and anatomical features of a large group of left and right predominant patients with TDP-43-C pathology. This will contribute towards more tailored criteria that specifically addresses the hallmark deficits caused by atrophy in the left or right ATL, as well as improve our understanding of the cognitive functions subserved by these lobes.

Variability in the clinical presentation of focal ATL neurodegeneration is likely linked to the extent and lateralization of atrophy within and between the temporal lobe, a structurally and functionally extremely heterogeneous portion of the cortex. Cellular, neurochemical, and pathological markers suggest the existence of at least seven distinct regions within the ATL (Ding, Van Hoesen, Cassell, & Poremba, 2009). Convergent evidence comes from in-vivo studies of structural and functional connectivity profiles. Based on whole brain functional connectivity patterns, Pascual and colleagues identified four major functional sub-regions within the ATL preferentially connected with the default-semantic network, paralimbic structures, visual networks, or auditory/somatosensory and language networks (Pascual et al., 2015). Similarly, structural connectivity analyses using diffusion tensor imaging support a structural parcellation within the ATL. For instance, Papinutto and colleagues demonstrated differential connectivity from a rostral region of the ATL to the orbitofrontal cortex (OFC), an anterior-lateral region to the occipital pole (OP), a ventro-lateral region to the middle temporal gyrus (MTG), a dorsal region to the superior temporal gyrus (STG), and two ventro-medial regions to the inferior temporal gyrus (ITG) and the fusiform gyrus (Fus) (Papinutto et al., 2016). Neuroimaging and neuropsychological investigations of the mosaic of functions subserved by the ATL has been hampered by the fact that it is highly susceptible to artefacts in fMRI (Visser, Jefferies, & Lambon Ralph, 2010) and rarely touched by strokes (but see (Tsapkini, Frangakis, & Hillis, 2011)). Instrumental to this end have been findings from neurodegenerative disease (Snowden et al., 1989), herpes simplex encephalitis (Kapur et al., 1994), temporal lobe epilepsy (Chabardès et al., 2005; Rice et al., 2018a; Rice et al., 2018b), and traumatic brain injury (Bigler, 2007). Overall, the ATL has been associated with language, in particular semantic knowledge (Binney, Embleton, Jefferies, Parker, & Lambon Ralph, 2010), but also socio-emotional cognition (Olson, Plotzker, & Ezzyat, 2007) and higher order visual, auditory and olfactory processes (Murray & Richmond, 2001). Notwithstanding this clear evidence of structural and functional subregions within the ATL, we currently lack an adequate description of regional atrophy distribution in TDP-43-C driven temporal variants of FTD. Moreover, while the longitudinal evolution of svPPA cases has been described (Brambati et al., 2009; Kumfor et al., 2016; Rohrer et al., 2008), atrophy progression of path-proven cases with asymmetric ATL involvement has never been compared.

In this study we investigate distribution and progression of TDP-43-C driven ATL neurodegeneration analyzing behavioral, imaging and clinical data in a sample of 30 pathology-proven cases of TDP-43-C having received a diagnosis of one of the temporal variants of FTD. First, we assessed the percentage of cases presenting with predominantly left vs. right atrophy applying an anatomical mask of the ATL. To investigate local atrophy distribution within each hemisphere, we then developed a novel parcellation of the ATL building on evidence from structural (Papinutto et al., 2016) and functional (Pascual et al., 2015) connectivity findings. While previous studies have focused on early involvement of medial temporal lobe substructures (i.e., hippocampi and amygdalae) in svPPA (Bocchetta et al. 2019), our parcellation is the first one to focus on the anterior temporal lobe and to probe possible differences along two axises: medial vs. lateral and anterior vs. posterior. Finally, we described the progression of atrophy within and outside the temporal lobe. We hypothesized that right-sided ATL degeneration is more common than previously thought and that atrophy distribution and progression would be similar, yet mirrored, in the two hemispheric variants.

## 2. Materials and methods

### 2.1 Participants

In this retrospective study, we included all patients in the database of the Memory and Aging Center at University of California, San Francisco (MAC, UCSF) and UCSF Neurodegenerative Disease Brain Bank (NDBB) that met the following inclusion criteria: (1) post-mortem neuropathological diagnosis of TDP-43-C, and (2) ante-mortem clinical diagnosis of one of the temporal variants of FTD as determined in the clinical records [i.e., semantic variant of PPA (svPPA (Gorno-tempini et al., 2011)), semantic dementia (SD (Neary, Snowden, Gustafson, Passant, Stuss, Black, Freedman, Kertesz, Robert, Albert, Boone, et al., 1998)), right variant of SD, temporal variant of FTD, or right variant of FTD - these latter being clinical diagnoses adopted at the MAC and not conforming to any consensus criteria]. Thirty-seven patients, all recruited between October 1, 1998 and January 31, 2014, met both our inclusion criteria. Of these patients, only five had a secondary contributing pathology: progressive supranuclear palsy (PSP) in two cases, primary lateral sclerosis (PLS) in the other two, and finally Alzheimer disease (AD) in one. Structural imaging was available for 30 of the 37 included patients. For 17 patients, three scans at least six months apart were available, allowing for additional longitudinal analyses in this subset. Table 1 and Results section 3.4 describe the demographic and neuropsychological profiles of our cohort. Twenty-five of the patients here analyzed have already been included in a previous publication (Spinelli et al., 2017). Of 437 cases in the NDBB (Kim et al., 2020), seven cases met criteria for one of the clinical syndromes but were excluded because they did not present with TDP-43-C pathology: three had Pick’s disease, two globular glial tauopathy, and two TDP- 43 type B with concomitant motor neuron disease and unclassifiable FTLD-tau pathology respectively. Additionally, two patients (90 and 79 years old respectively at the time of death) showed TDP-43-C pathology at autopsy but did not meet clinical criteria for any FTD clinical syndrome, rather for mild cognitive impairment (MCI). These two cases might be seen as prodromal cases of FTD, raising interesting questions on which clinical and cognitive features would left- and right- sided temporal FTD have at very early stages. However, an in-depth description of these cases, hampered by the minimal clinical data available (i.e., they were not followed clinically and thus did not undergo the full UCSF MAC testing protocol), would go beyond the scope of the present manuscript. The study was approved by the UCSF Committee on Human Research and all subjects provided written informed consent.

**Table 1.**
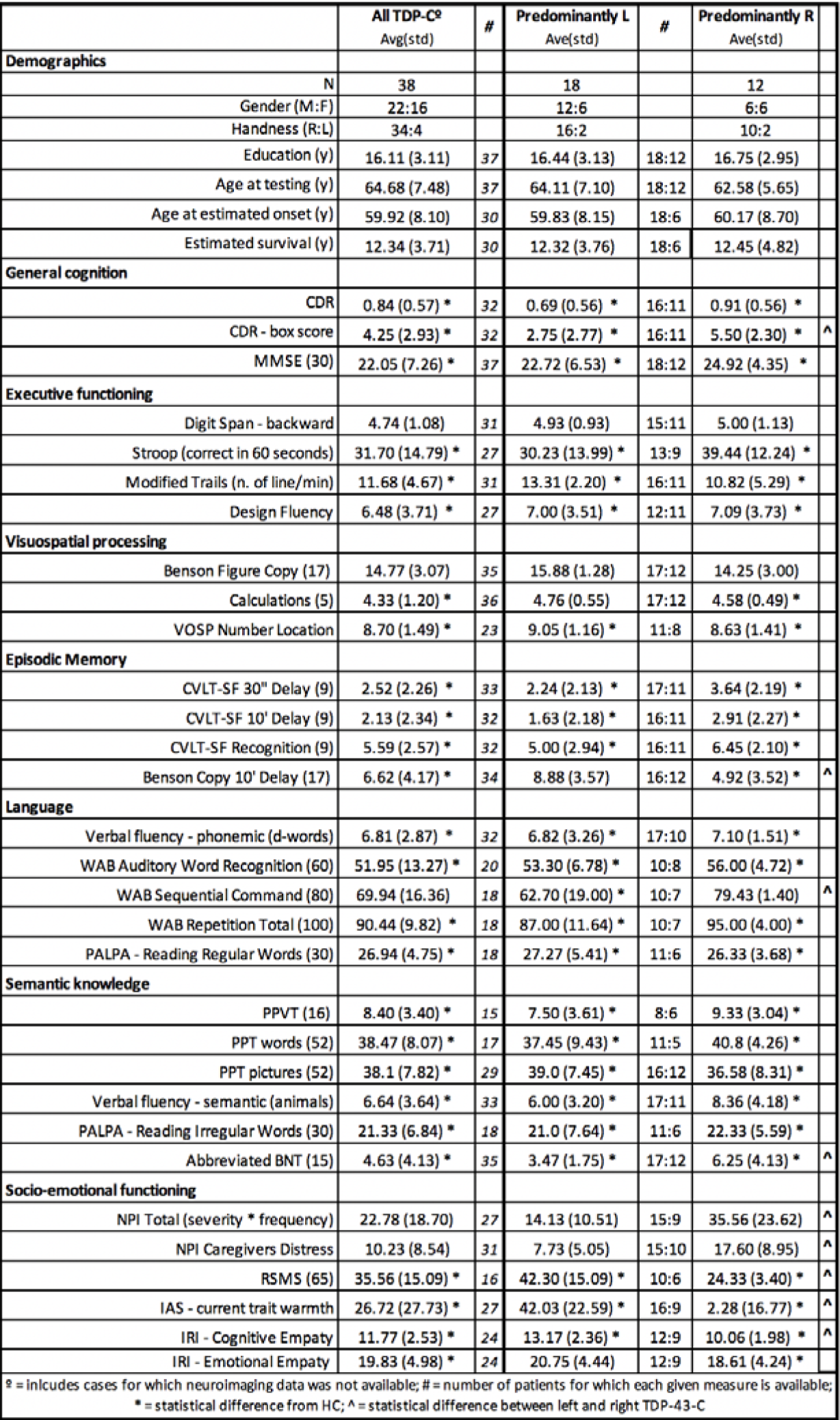
Demographic and neuropsychological characteristics of the participants. Scores shown are mean (standard deviation), with asterisks (*) indicate values significantly different from controls, and carets (^) statistical differences between predominantly left and right TDP-43-C cases. CDR = Clinical Dementia Rating; MMSE = Mini Mental State Exam; VOSP = Visual Object and Space Perception Battery; CVLT-SF = California Verbal Learning Test-Short Form; WAB = Western Aphasia Battery; PALPA = Psycholinguistic Assessments of Language Processing Abilities; PPVT = Peabody Picture Vocabulary; PPT = Pyramid and Palms Tree; BNT = Boston Naming Test; NPI = Neuropsychiatric Inventory; RSMS = Revised Self-monitoring Scale; IAS = Interpersonal Adjective Scales; IRI = Interpersonal Reactivity Index.

### 2.2 Neuropathological, genetic, and neuropsychological assessment

Thirty-one autopsies were performed at UCSF, and six were performed at the University of Pennsylvania. Primary and secondary pathological changes were established by the pathologist based on consensus criteria (Kovacs et al., 2016; Mackenzie et al., 2011; McKeith et al., 1996; Montine et al., 2012).

For 30 out of the 37 participants who donated their brains, blood samples were available. Following previously described protocols (Li et al., 2014; Moreno et al., 2015), we screened for known pathogenic mutations in the following genes: GRN, MAPT, TARDBP, C9ORF72, APP, PSEN1, PSEN2, FUS. Apolipoprotein E (APOE) and MAPT H1/H2 haplotypes were also assessed. Twenty-five of these 30 cases also had available structural imaging.

Clinical diagnosis was based on a detailed medical history, comprehensive neurological exam, and standardized neuropsychological and language evaluations (Kramer et al., 2003). At the time of recruitment, the consensus criteria used for the evaluation (i.e., guiding the choice of tests and the final clinical diagnosis) were those of Neary et al., 1998. According to those criteria, out of the 30 cases for which both behavioral and imaging data was available, 28 met criteria for SD and 2 cases met criteria for FTD. If the current criteria for svPPA were to be applied retrospectively (Gorno-Tempini et al., 2011), only 19 cases would meet both root criteria for PPA (prominent, early language complaints) and specific svPPA criteria.

Demographic and neuropsychological characteristics of the patients included in the study are shown in Table 1. Two-sample t-tests (two-tailed distributions, significance threshold set at p<0.05) were used to statistically assess group differences between healthy controls published data (Gorno-Tempini et al., 2004; Watson et al., 2018) and our TDP-43-C patients, considered as an undifferentiated cohort as well as split in two groups following the neuroimaging results described later (i.e., left-predominant vs. right-predominant atrophy pattern). Similarly, we directly compared left-predominant and right-predominant groups to investigate atrophy lateralization effects on the neuropsychological profile.

### 2.3 Neuroimaging protocols

For the neuroimaging analysis, a set of thirty healthy controls (HC, 18 females, mean age 65.1±8.7) matched with patients for age, gender, and scanner type was included from the MAC UCSF Hillbloom healthy aging cohort. 3D T1-weighted images with a magnetization-prepared rapid gradient echo sequence (MPRAGE) were obtained from patients and HC using either a 1.5 (n=25 in both cohorts), 3 (n=3 in both cohorts), or 4 (n=2 in both cohorts) Tesla scanners with the following parameters. For 1.5T images: Siemens Magnetom VISION system (Siemens, Iselin, NJ), standard quadrature head coil, 8-channel, 164 coronal slices; repetition time (TR) = 10 ms; echo time (TE) = 4 ms; inversion time (TI) = 300 ms; flip angle = 15°; field of view (FOV) = 256 × 256 mm2; matrix 256 × 256; voxel size 1.0 × 1.5×1.0 mm2; s. For 3T images: Trio Siemens, 8-channel receive head coil, 160 sagittal slices, TR=2300 ms, TE =2.98 ms, TI = 900 ms, flip angle 9o, FOV = 256×256 mm, matrix size = 256×240, voxel size 1.0 × 1.0×1.0 mm3;. For 4T images: Bruker/Siemens, single housing birdcage transmit and 8-channel receive coil, 157 sagittal slices; TR = 2300 ms; TE = 3 ms; TI = 950 ms; flip angle = 7°; FOV = 256×256 mm2; matrix 256 × 256; voxel size 1.0 × 1.0×1.0 mm2.

### 2.4 ATL parcellation

We investigated regional ATL susceptibility to TDP-43-C driven neurodegeneration with a trade-off between anatomical and cytoarchitectonic specificity. Given the spatial resolution of our T1 images, and the spread of atrophy observed even in the earliest cases, the fine-grained partition provided by cytoarchitectonic and chemoarchitectonic markers could not be used directly (Ding et al., 2009). Rather, we parcellated both ATLs into four regions of interest based on previous findings of dissociable functional and structural connectivity profiles which suggested four-to-six main subdivisions (Papinutto et al., 2016; Pascual et al., 2015). In particular, the structural partition suggested by Papinutto and colleagues, based on white matter connectivity of the ATL to cortical areas, was used to draw, in the MNI space, four ROIs in each hemisphere. These ROIs correspond to the anterior and posterior portions of the medial and lateral ATL respectively, thus covering both paralimbic and neocortical structures (see Figure 4A). In detail, they isolate:

**Figure 1.**
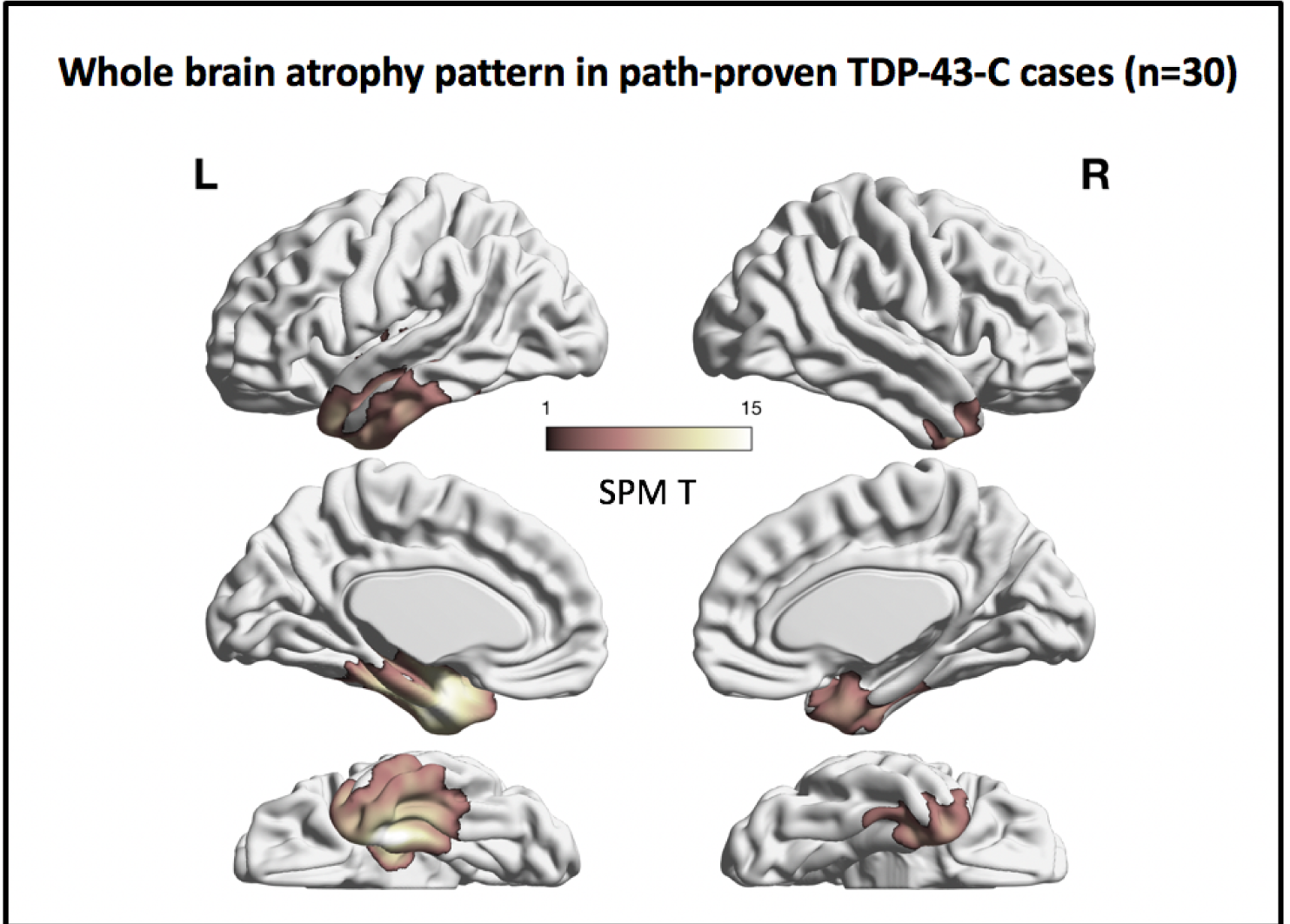
Atrophy distribution in 30 path-proven TDP-43-C cases. Render illustrating the results of the voxel-based morphometry (VBM) analysis identifying regions of grey matter volume loss in our sample of TDP-43-C patients (n=30) relative to age-matched healthy controls. Statistical maps are thresholded at p<0.05 corrected at the cluster level for family-wise error (FWE), with a cluster extended threshold of 100 voxels. Covariates: age, gender, handiness, GM, and scanner type. Details of the significant clusters are in Table 2.

We then investigated subject-specific atrophy asymmetry in a data-driven fashion. Relying on anatomical masks of the two ATLs (Figure 2A), we computed an index of atrophy lateralization (see Methods) an automatically classified each patient as left-predominant or right-predominant. Eighteen cases were found to have left-lateralized atrophy (60%), with lateralization values ranging from a maximum value of 0.22 to a minimum of 0.01 (mean: 0.15, std: 0.06, median: 0.18). In twelve cases, a right-lateralized atrophy pattern was detected (40%), with lateralization values ranging from a maximum value of −0.24 to a minimum of −0.07 (mean: −0.14, std: 0.05, median: −0.13). Figure 2b illustrates the distribution of volume loss across left and right ATLs in the whole sampl (n=30), as well as separately for predominantly left (n=18) and right cases (n=12). More variability can be appreciated on the right hemisphere, in particular in predominantly left cases. The within-group distribution of the lateralization index is shown in Figure 2c: atrophy lateralization describes a continuum, with slightly more variability in the predominantly right group.

**Figure 2.**
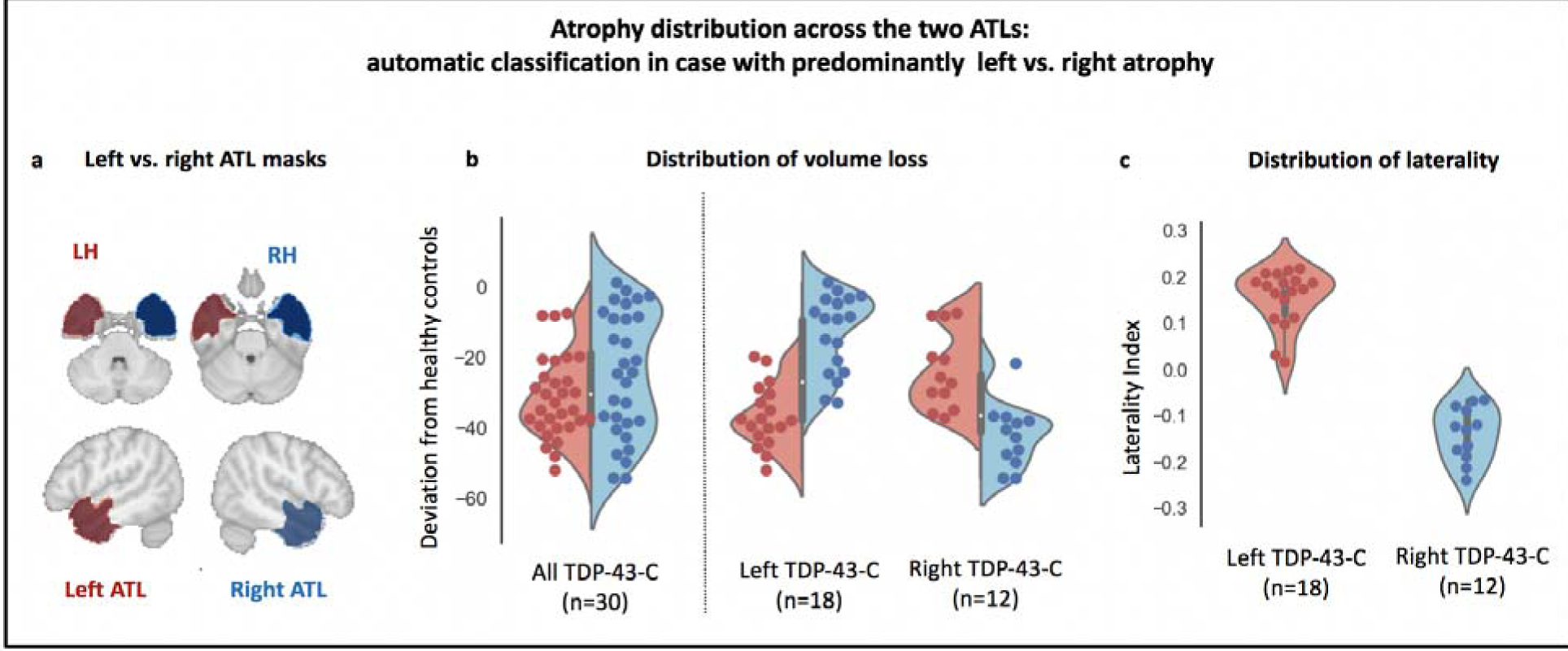
Atrophy distribution between the ATLs: automatic classification in left-predominant vs. right-predominant cases. A) Mask of the anterior temporal lobe (ATL) used to assess atrophy lateralization index. B) Violin plots illustrating subjects’ mean percentage of volume loss in the left (red) and right (blue) ATL for the undifferentiated cohort of TDP-43-C cases (n=30), for cases with a positive atrophy lateralization index (i.e., atrophy predominantly affecting the left ATL (n=18), and for cases with a negative atrophy lateralization index (i.e., right-predominant ATL atrophy (n=12). C) Violin plot illustrating subjects’ laterality index in predominantly left (red, n=18) and predominantly right (blue, n=12) cases.

**Figure 3.**
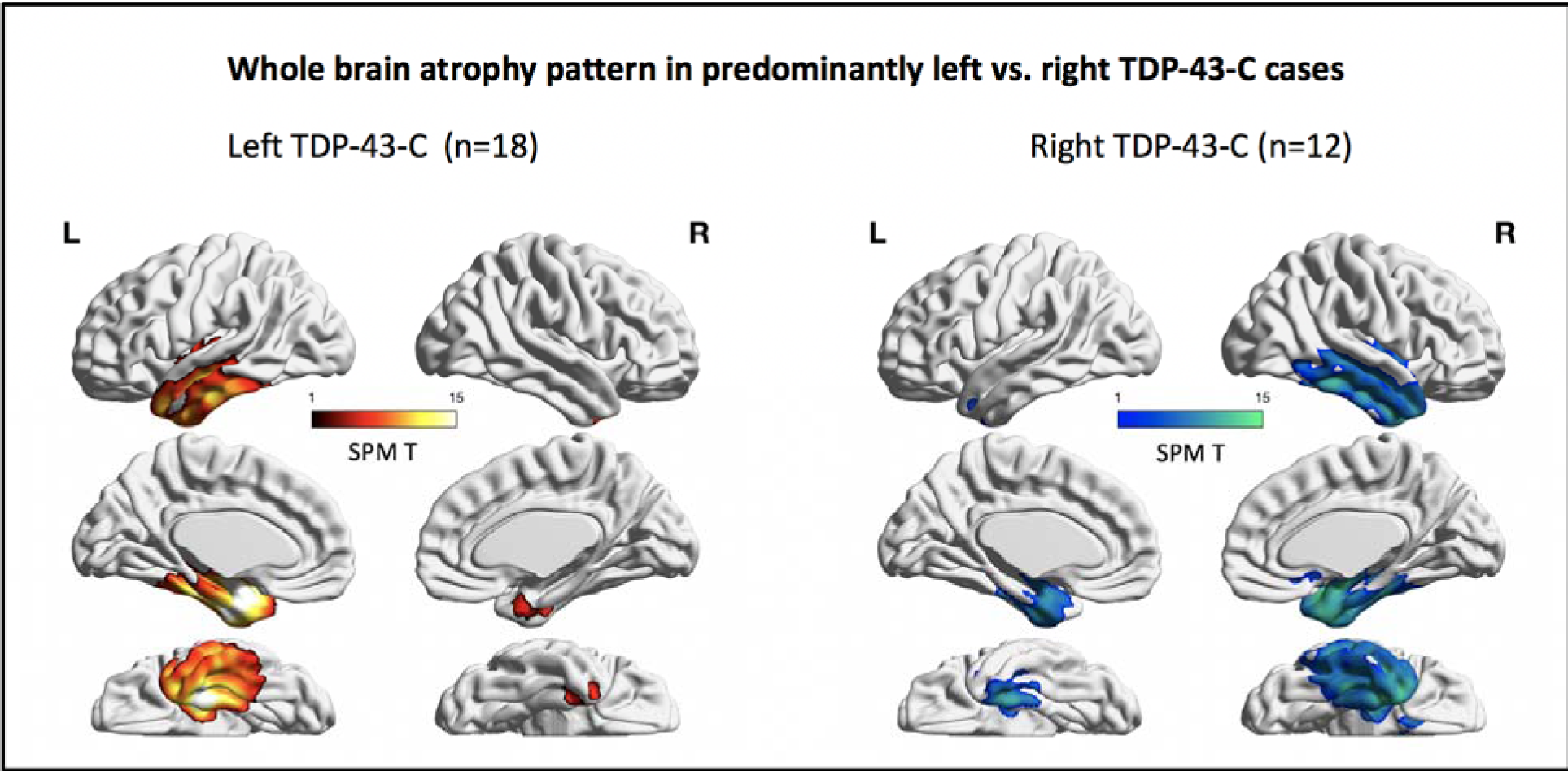
Voxel-based morphometry (VBM) for left-predominant (n=18, red) and right-predominant (n=12, blue) cases. Maps are thresholded at p<0.05 corrected at the cluster level for family-wise error (FWE), with a cluster extended threshold of 100 voxels. Covariates: age, gender, handiness, GM, and scanner type. Details of the significant clusters are in Table 2.

**Figure 4.**
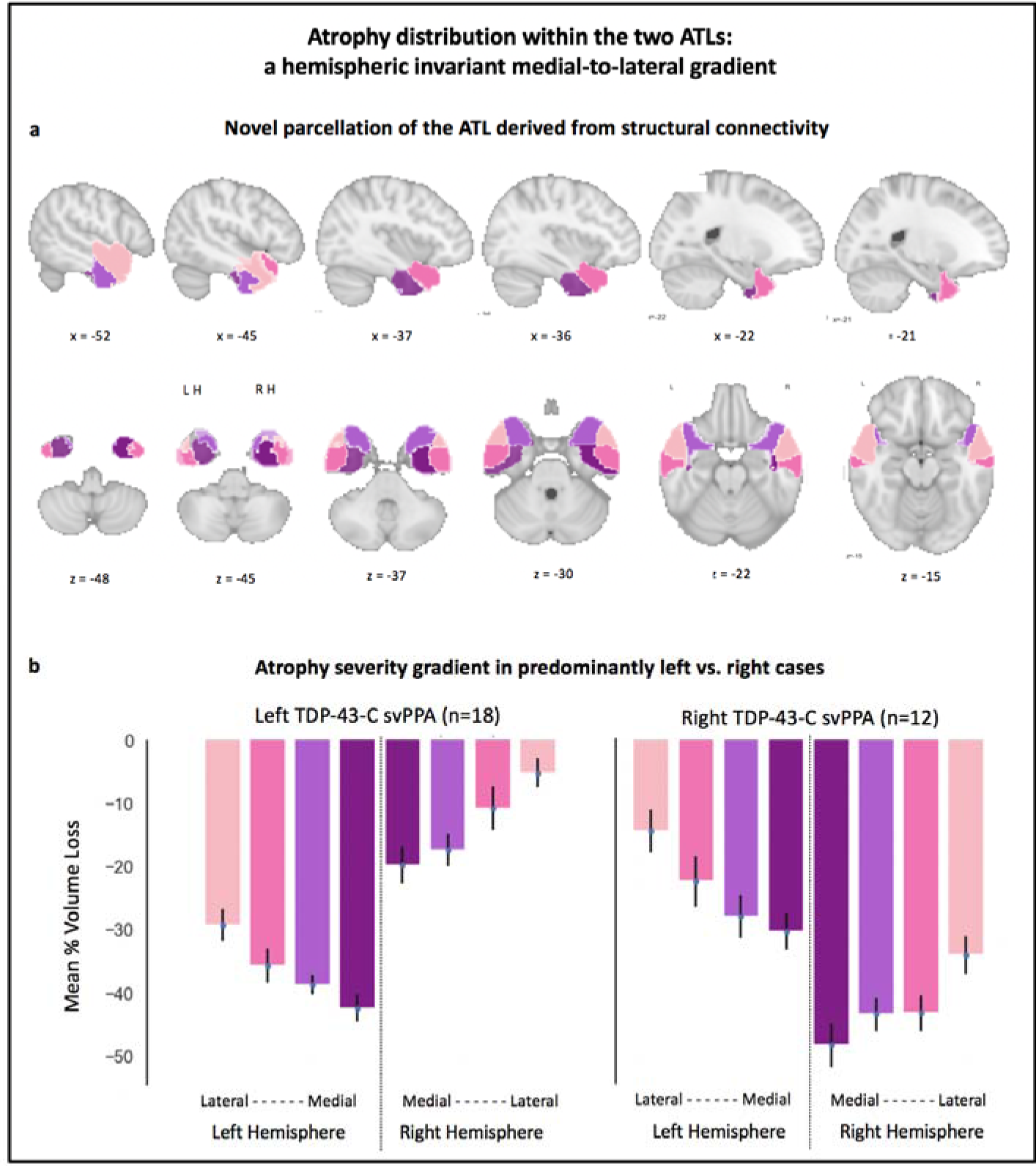
Atrophy distribution within ATLs: hemispheric invariant medial-to-lateral gradient. A) Regions of interest (ROIs) drawn to parcellate the anterior temporal lobe in regions with known differences in cytoarchitecture, as well as structural and functional connectivity. B) Mean percentage of volume loss in the 4 bilateral ROIs for left-predominant (left, n=18) and right-predominant (right, n=12) TDP-43-C svPPA. Error bars represent standard error of the mean (SEM).

⍰ an inferior-medial region, in Papinutto (2016) preferentially connected with inferior occipital pole and inferior temporal gyrus, encompassing areas 35 and 36 as described in Ding (2009);

⍰ a superior-medial region, in Papinutto (2016) preferentially connected with orbitofrontal cortex, and including areas TG and TE (Ding et al., 2009);

⍰ an inferior-lateral region, in Papinutto (2016) preferentially connected with middle temporal gyrus, and including areas EC and TI (Ding et al., 2009);

⍰ a superior-lateral region, in Papinutto (2016) preferentially connected with superior occipital pole and superior temporal gyrus, and encompassing area TA (2009).

In absence of an anatomical landmark, a sagittal plane at x=42 was used to define the boundary between two of the ROIs derived from Papinutto (2016): the inferior-medial region (preferentially connected with inferior temporal and occipital lobes) and the superior-lateral one (preferentially connected with superior temporal and occipital lobes). Finally, the sum of the four ipsilateral ROIs lead to the development of two masks for left vs. right ATLs, used to compute an index of atrophy lateralization (see below). This novel parcellation of the ATL, the first one allowing the appreciation of medial vs. lateral and anterior vs. posterior effects, has never been used in a sample of path-proven TDP-43-C cases yet we have recently used it to describe the longitudinal evolution of a single case of svPPA in Vonk et al., 2019.

### 2.5 MRI cross-sectional analyses

Cross-sectional voxel-based morphometry (VBM) analysis of the structural images was conducted to assess volume differences across cohorts, as later described. T1-weighted images undergone bias field correction using N3 algorithm, and segmented into gray matter (GM), white matter (WM) and cerebrospinal fluid (CSF) using the unified segmentation (Ashburner & Friston, 2005) in Statistical Parametric Mapping (SPM12) (Wellcome Trust Center for Neuroimaging, London, UK; http://www.fil.ion.ucl.ac.uk/spm/software/spm12/) running on Matlab R2013a (MathWorks). A custom group template was generated from the segmented gray and white matter tissues and cerebrospinal fluid by non-linear registration template generation using Large Deformation Diffeomorphic Metric Mapping framework (Ashburner & Friston, 2011). Segmented images in the native space were then registered to the custom template and modulated by the Jacobian determinant to preserve the relative GM volume. For statistical purposes, linear and non-linear transformations between the group template space and International Consortium of Brain Mapping (ICBM, Fonov et al. 2009) were applied. All steps of the transformation were carefully inspected from the native space to the group template. Finally, images were smoothed for statistical analysis (8 mm full-width at half-maximum [FWHM] Gaussian kernel). Whole-brain differences in GM volume were investigated using a general linear model including as covariates those variables known to determine gross anatomical differences (i.e., age, gender, handiness), scanner type and a proxy for disease severity (i.e., total GM volume). The best available proxy was total GM volume, a continuous, global variable objectively measurable in all subjects at the first visit. We lacked in fact an accurate estimation of disease onset, and no single neuropsychological score could be used: symptoms vary considerably, not all neuropsychological measures are available for all subjects, and most summary scores of general cognitive status are not continuous and/or have very little variability (Good et al. 2001, Canu et al. 2020). It should be noted that estimations of time-sensitive variations (e.g., age-related changes) based on GM-adjusted data generalize better to new samples, suggesting that GM correction should be preferred to other measures when lacking additional information and/or specific hypotheses (Peelle, et al., 2012). To assess cross-sectional whole-brain differences in GM volume, three group comparisons were performed: HC vs. all TDP-43-C cases, HC vs. predominantly left TDP-43-C cases, HC vs. predominantly right TDP-43-C cases. Whole-brain between-group statistically significant differences in GM volume were explored at p<0.05 corrected at the cluster level for family-wise error (FWE), with a cluster extended threshold of 100 voxels.

We further characterized the pattern of atrophy between and within temporal lobes by using our ad-hoc parcellation, grounded in structural evidence of ATL heterogeneity. First, for each subject, we computed the average GM value in each ROI. This value was scaled by the overall GM (to control for global differences in head size and disease severity, in keeping with the whole-brain VBM analyses) and normalized dividing it by the average volume in HC to express distance from normal values. We then calculated a laterality index as the ratio between the difference in volume between the two hemispheres and their sum, i.e.: *(Av_Right - Av_Left) / (Av_Right + Av_Left)*, where (as described above) the masks used to compute *Av_Right and Av_Left* were given by the sum of the four ROIs on the right and left hemisphere respectively. A positive index thus indicates predominantly left atrophy, while a negative one denotes greater atrophy in the right ATL. This index allowed for a data-driven classification of all TDP-43-C cases as either predominantly left or predominantly right (fixing the threshold for classification at 0). It should be noted that this measure of atrophy asymmetry is relative to the single subject, i.e., established for each individual case, not resulting from a comparison to the group. The label assigned identifies the hemisphere most affected intra-individually; it does not preclude the possibility that the non-predominantly affected hemisphere might also be severely affected, when compared to controls or other patients.

We then visualized and statistically compared volume loss across subjects, ROIs, and hemispheres, aiming to assess possible modulations of the within-ATL atrophy distribution. Atrophy scores (i.e., 8 for each participant, corresponding to the averaged, normalized, and total GM corrected volume in our ROIs) were entered in two linear mixed effect models. The first one included fixed effects for atrophy lateralization (i.e., binary classification in predominantly left vs. right cases, as defined by our laterality index), hemisphere (i.e., left vs. right ATL), ROIs (i.e., 4 regions from medial to lateral), and a random by-participant intercept. This model allowed for all possible two- and three-ways interactions between all main effects (e.g., is the across-ROIs distribution of atrophy modulated by the hemisphere considered? and/or by the side of prevalent atrophy?). As we observed a lack of significant two- and three- way interactions between the main effect of ROIs (i.e., the medial-to-lateral gradient) and the other two main effects (i.e., atrophy lateralization and hemisphere), we fit a second, restricted model. This model considered only the main effects (i.e., atrophy lateralization, hemisphere, ROIs) and the possible interaction between atrophy lateralization and hemisphere, thus excluding possible two- and three- ways interaction with ROIs. It should be noted that this latter interaction is trivial and expected since patients’ atrophy lateralization was established based on the hemisphere being most affected. We then statistically compared the results of two models via a likelihood ratio test. A similar performance between the full and restricted models would strongly suggest that the variance in the data is explained by the main effects (i.e., atrophy lateralization, hemisphere, and ROIs) and by the expected interaction between atrophy lateralization and hemisphere, with no modulation of the ROIs effect by neither atrophy lateralization nor hemisphere. Additionally, we explored the possibility that supplementary information could be provided by considering atrophy lateralization as a continuous measure instead of the categorical grouping in predominantly left vs. right cases. We re-fitted (and compared) the two models replacing the binary classification (i.e., predominantly left vs. predominantly right) with the raw index of lateralization.

Finally, we correlated the average volume loss in our eight ROIs with two tests that measure key neuropsychological features of left and right temporal variants of FTD respectively (Binney et al., 2016): PALPA – exception word reading (a measure of surface dyslexia caused by verbal semantic deficits) and the Neuropsychiatric Inventory (a measure of behavioral symptoms).

### 2.6 MRI longitudinal analyses

All T1-weighted images underwent bias field correction using N3 algorithm, and then segmented using SPM12 unified segmentation (Ashburner & Friston, 2005). An intra-subject template was created by non-linear diffeomorphic and rigid-body registration proposed by the symmetric diffeomorphic registration for longitudinal MRI framework (Ashburner & Ridgway, 2013). The intra-subject template was segmented using SPM12’s unified segmentation. A within-subject modulation was applied by multiplying the timepoints’ jacobian with the intra-subject averaged tissues (Ziegler et al., 2015). A custom group template was generated from the within-subject template gray and white matter tissues and cerebrospinal fluid by non-linear registration template generation using Large Deformation Diffeomorphic Metric Mapping framework (Ashburner & Friston, 2011). In the intra-subject template, we capture the volume variation between two timepoints by subtracting the timepoint’s jacobian. We estimate the rate of change by dividing the variation of tissue density with respect of the time difference between two time-points. This rate of change between two time-points is then pushed to the group template using the subject composition of transformations field. For statistical purposes, linear and non-linear transformations between the group template space and ICBM (Fonov et al. 2009) were applied. Every step of the transformation was carefully inspected from the native space to the group template. Finally, for statistical purposes, images were smoothed with a Gaussian kernel (4 mm FWHM).

Whole-brain differences of gray matter changes were investigated using a general linear model including age, gender, handiness, and scanner type as covariates. Two group comparisons were performed: HCs vs. predominantly left TDP-43-C cases (n=13), HCs vs. predominantly right TDP-43-C cases (n=4). We repeated the process twice to compute change maps between time point 1 and time point 2, as well as between time point 2 and time point 3. Results are shown with a threshold of significance set at p<0.05 corrected for family-wise error (FWE), with a cluster extended threshold of 100 voxels. Subsequently, a more liberal threshold at p<0.001, uncorrected, was explored to avoid false negatives that can occur in small groups’ sample size. Overall, it should be noted that given the extremely small sample of predominantly right TDP-43-C cases with longitudinal scans available (n=4), the results referring to this subset of our original cohort are purely descriptive.

## 3. Results

### 3.1 Atrophy distribution between the ATLs: imaging-based classification of left vs. right variants

We sought to characterize the pattern of atrophy distribution between the two ATLs in a data-driven, imaging-based way. The undifferentiated cohort of TDP-43-C cases presented with the expected pattern of atrophy, involving bilateral ATLs, but with greater volume loss in the left hemisphere (Figure 1, Table 2). Consistent with previous studies in svPPA (Chan et al., 2001; Galton et al., 2001; Mummery et al., 2000; Nestor, Fryer, & Hodges, 2006), the most severely affected regions of the left temporal lobe included the temporal pole (BA 38), the fusiform (BA 20) and lingual (BA 37) gyri. In the right hemisphere, significant volume loss was found in the temporal pole (BA 38) and inferior (BA 20) temporal gyri. In both hemisphere amygdala, hippocampus and parahippocampal regions were severely affected, with an anterior-to-posterior gradient (Binney et al., 2016; Chan et al., 2001).

**Table 2.** Results of the voxel-based morphometry (VBM) analysis. Maps were thresholded at p<0.05 corrected at the cluster level for family-wise error (FWE), with a cluster extended threshold of 100 voxels. Covariates: age, gender, handiness, GM, and scanner type. Render of the significant clusters are displayed in Figure 1 and 3. MNI [x,y,z] = coordinates in MNI space; p-value = FWE corrected p-value at the cluster level; Ke = number of voxels; t-value = maximum T statistic at each peak.

| Table 2. Results of the voxel-based morphometry (VBM) analysis |  |  |  |  |
| --- | --- | --- | --- | --- |
|  | Local Maxima |  |  |  |
|  | MNI [x,y,z] | p-value | Ke | t-value |
| *Path-proven TDP-43-C case (n=30)* |  |  |  |  |
| left parahippocampal gyrus (BA36) | -26 -14 -28 | <0.001 | 25394 | 17.22 |
| right temporopolar area (BA38) | 30 8 -34 | <0.001 | 6133 | 8.68 |
| *Left-predominant Cases (n=18)* |  |  |  |  |
| left parahippocampal gyrus (BA36) | -26 -14 -27 | <0.001 | 30084 | 19.69 |
| right temporopolar area (BA38) | 30 8 -36 | <0.001 | 887 | 7.11 |
| *Rght-predominant Cases (n=12)* |  |  |  |  |
| right temporopolar area (BA38) | 39 3 -16 | <0.001 | 24746 | 16.3 |
| left parahippocampal gyrus (BA36) | -26 -14 -28 | <0.001 | 5164 | 11.4 |
| right orbitofrontal (BA11) | 9 18 -16 | <0.001 | 577 | 7.73 |

To assess potential differences outside the temporal lobes, whole-brain VBM results for the two groups are shown in Figure 3. Left-predominant cases show significant gray matter volume loss in the left temporal pole (BA 38), left fusiform (BA 20) and left lingual (BA 37) gyri, as well as in the right temporal pole. In right-predominant cases, we observed significant atrophy not only in the right temporal pole, right fusiform and right lingual gyri, but also in the left temporal pole and left inferior frontal gyrus. These results are in line with previous studies comparing left and right temporal variants of FTD (Binney et al., 2016; Brambati et al., 2009; Kumfor et al., 2016; Rogalski et al., 2014). It should be noted that direct comparison of the two groups is hampered by an inevitable difference in sample size (18 left-predominant cases vs. 12 right-predominant ones).

### 3.2 Atrophy distribution within ATLs: hemispheric invariant medial-to-lateral gradient

Given the known anatomical heterogeneity of the temporal pole, we aimed to describe and compare atrophy distribution within the two ATLs. We examined regional differences thanks to a novel parcellation of the ATL based on recent findings on its structural connectivity profile (Figure 4A). In both hemispheres, and irrespective of atrophy lateralization, medial regions of the ATLs appeared to be more affected by volume loss than lateral ones (Figure 4B). A linear mixed effect model revealed a significant effect of atrophy lateralization (left-predominant vs. right-predominant cases, z= 3.048, p<0.002) and hemisphere (left vs. right ATL, z= 10.33, p<0.001). Moreover, all four ROIs were significantly different in volume (all p <.05), except the two most medial ROIs, which were only marginally significantly different (p= 0.08). Unsurprisingly, given how atrophy lateralization was computed, there was a significant two-way interaction between atrophy lateralization and hemisphere (z=- 11.7, p<0.001). Crucially, none of the other two- and three- way interactions were significant, indicating that atrophy lateralization did not modulate across-ROIs atrophy distribution, nor did hemisphere, or their combination. This null finding suggests that the within-ATL pattern of atrophy (i.e., the medial-to-lateral gradient) is not modulated by either atrophy lateralization (i.e., atrophy predominantly affecting the left or right ATL) or hemisphere (i.e., whether considering the left vs. right ATL). To corroborate this observation, we ran a second, restricted, model that did not include interaction terms for the two- and three- way interactions involving ROIs. This model confirmed our previous results, indicating a main effect of atrophy lateralization (left-predominant vs. right-predominant cases, z= 3.8, p<0.001), hemisphere (left vs. right ATL, z= 20.89, p<0.001), and ROIs (all ps <.05), and a significant two-way interaction between atrophy lateralization and hemisphere (z=-23.72, p<0.001). Critically, the full and restricted models did not statistically differ from each other (Chisq = 5.07, p =. 82), supporting the idea that the variance in the data is explained by the main effects (i.e., atrophy lateralization, hemisphere, and ROIs) and by the expected interaction between atrophy lateralization and hemisphere, with no modulation of the ROI effect by atrophy lateralization or hemisphere. Finally, the same main effects, and two- / three- ways interactions were found when the two models were re-fitted entering atrophy lateralization as a continuous variable (i.e., the raw lateralization index), instead of a binary classification. Even with these new models the likelihood test comparing the full and restricted versions failed to reach significance. Overall, these findings support the conclusion that there is no interaction between ROIs (i.e., from medial to lateral) and either hemisphere (left vs. right ATL) or atrophy lateralization (i.e., cases with predominantly left vs. right atrophy).

### 3.3 Atrophy longitudinal progression: a mirror evolution

After establishing the pattern of atrophy distribution between and within the two ATLs, we compared longitudinal changes in left- and right-predominant TDP-43-C cases. Between time point 1 and time point 2, left-predominant patients lost volume in right anterior temporal and left posterior temporal lobes (Figure 5A, in red). Similarly, right-predominant cases showed increased atrophy in left anterior temporal and right posterior temporal lobes (Figure 5A, in blue). Comparing time point 2 with time point 3, both variants show mirrored spreading of atrophy to more posterior regions of both temporal lobes, as well as more involvement of the ipsilateral orbitofrontal cortex (Figure 5B). These results, in line with previous findings (Brambati et al., 2009; Kumfor et al., 2016), indicate a progressive merging of the two variants to a common profile of bilateral temporal atrophy. It should be noted that the right-predominant cases included in these analyses are only four and that when the whole longitudinal sample is considered (n=17), the pattern of change was driven by left-predominant cases progressing contralaterally to atrophy on the right, as previously reported (Rohrer et al., 2008). Given the size of the available sample, the current analysis can only illustrate how atrophy progression in TDP-43-C path-proven cases conforms with previous findings on temporal FTDs, while calling for future studies, in larger samples, including both predominantly left and right cases.

**Figure 5.**
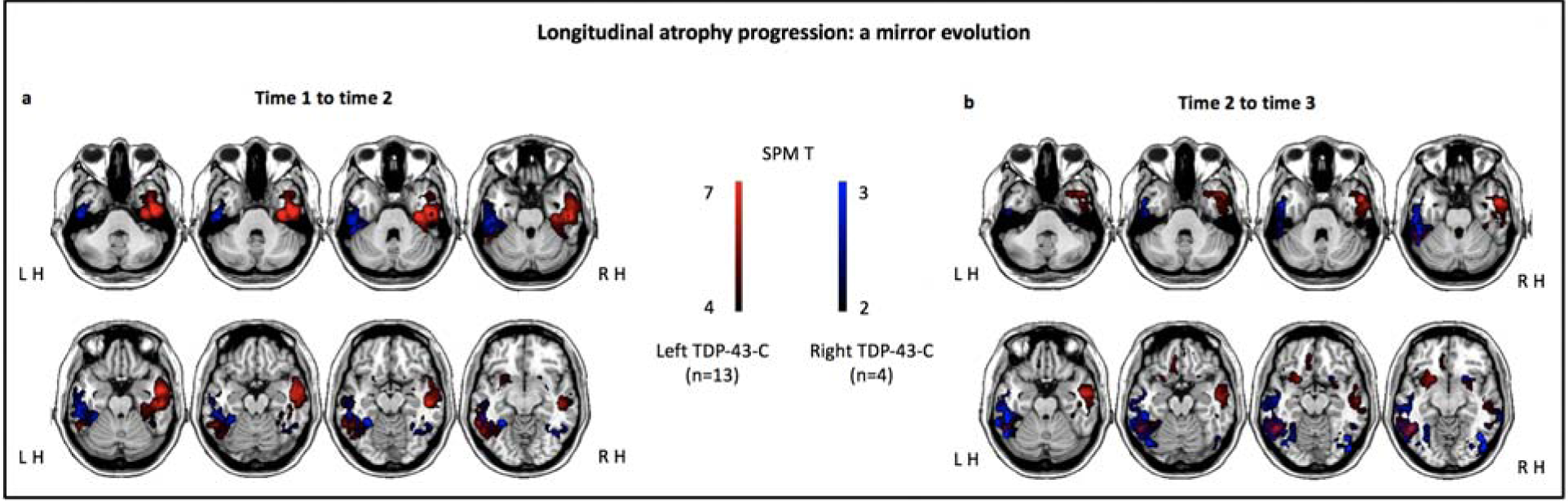
Longitudinal atrophy progression. Overlay of the change maps for left-predominant (in red, n=13) and right-predominant (in blue, n=4) TDP-43-C cases between time 1 and time 2 (A) and between time 2 and time 3 (B). Given the small sample size, maps are thresholded at p < 0.001 uncorrected. Covariates: age, gender, handiness, and scanner type.

### 3.4 Lateralization of ATL neurodegeneration drives two clinical phenotypes

To better characterize TDP-43-C driven ATL neurodegeneration, we first describe the demographic, genetic and neuropsychological profile of all patients for which behavioral data is available. We then focus on those included in the cross-sectional imaging portion of the study, which allows us to illustrate the phenotypical differences between left- and right-predominant TDP-43-C cases. It should be noted that given the retrospective and path-confirmed nature of the study, not all measures are available for all subjects. Table 1 reports all data, as well as the number of subjects for which a given score is available.

As per selection criteria, all 37 patients included in the study (16 females, four left-handed, mean (SD) age = 64.68 (± 7.48) years, mean (SD) education = 16.11 (± 3.11) years) had received a diagnosis falling within the umbrella of temporal variants of FTD. At the time of their first evaluations (between 1998 and 2012), all patients met Neary criteria (Neary, Snowden, Gustafson, Passant, Stuss, Black, Freedman, Kertesz, Robert, Albert, & Boone, 1998). Specifically, they all meet criteria for semantic variant of FTD (SD in Neary et al., 1998), except for two who met criteria for behavioral variant (FTD in Neary et al., 1998, both right-predominant according to our atrophy lateralization index). It should be noted that nine cases could be diagnosed with either SD or FTD as they fulfille both criteria (1 left-predominant and 8 right-predominant) and that all but one of the right-predominant cases would fail to meet current criteria for svPPA, as not meeting general PPA criteria first (Gorno-tempini et al., 2011).

Within the pool of 30 patients for which genetic data were available, no patient showed a known pathogenic variant in an FTLD-associated gene. The ApoE4 allele was present in three patients (one left-predominant, two right-predominant). The H1/H1 MAPT haplotype was found in 56% of the cases, with no significant differences between predominantly left vs. right cases (left-predominant = 7:7, right-predominant = 9:4, χ^2^ =1.03, p=0.31).

The neuropsychological characteristics of the undifferentiated cohort of TDP-43-C cases are consistent with the expected profile for temporal FTD and summarized in Table 1. Significant differences from healthy controls indicate semantic deficits and impaired comprehension with no sign of apraxia of speech or dysarthria. In line with previous evidence, the same overall pattern is observed when left-predominant and right-predominant cases are considered separately, yet interesting distinctions can also be appreciated (Binney et al., 2016; Lambon Ralph et al., 2001; M. Mendez, Kremen, Tsai, & Shapira, 2010; Mion et al., 2010). The two groups appear to be well matched in terms of education, age at testing, and age at onset. Global cognitive performance was comparable, with no significant difference in MMSE and CDR global score. Left-predominant patients presented with greater impairment in object naming (short BNT, t=-2.18, p = 0.04) and following sequential commands (WAB Sequential Commands, t=-2.63, p = 0.02). Right-predominant cases had significant more behavioral disturbances (CDR-box scores, t=-2.67, p = 0.01 and NPI total, t=-2.43, p = 0.03), less socioemotional sensitivity (RSMS, t=3.36, p = 0.006), less interpersonal warmth (IAS, t=-4.78, p = 0.0001), less cognitive empathy (IRI, t=3.12, p = 0.005), and worse visuo-spatial performance (modified Rey figure delayed copy, t=2.81, p = 0.009).

Finally, we investigated the relation between atrophy in our novel ROIs and key neuropsychological features (see Figure 6). Reading performance in the irregular words subtest of the PALPA, a proxy for verbal semantic efficiency, correlated with volume loss in all left hemisphere ROIs (and none of the right), with the strongest correlation being detected with the most lateral ROI (r = 0.69, p = 0.002). Psychiatric symptoms, detected with the NPI and denoting behavioral changes, showed significant correlations with all right hemisphere ROIs (and none of the left), with the strongest correlation being detected with the most medial ROI (r = −0.62, p = 0.001).

**Figure 6.**
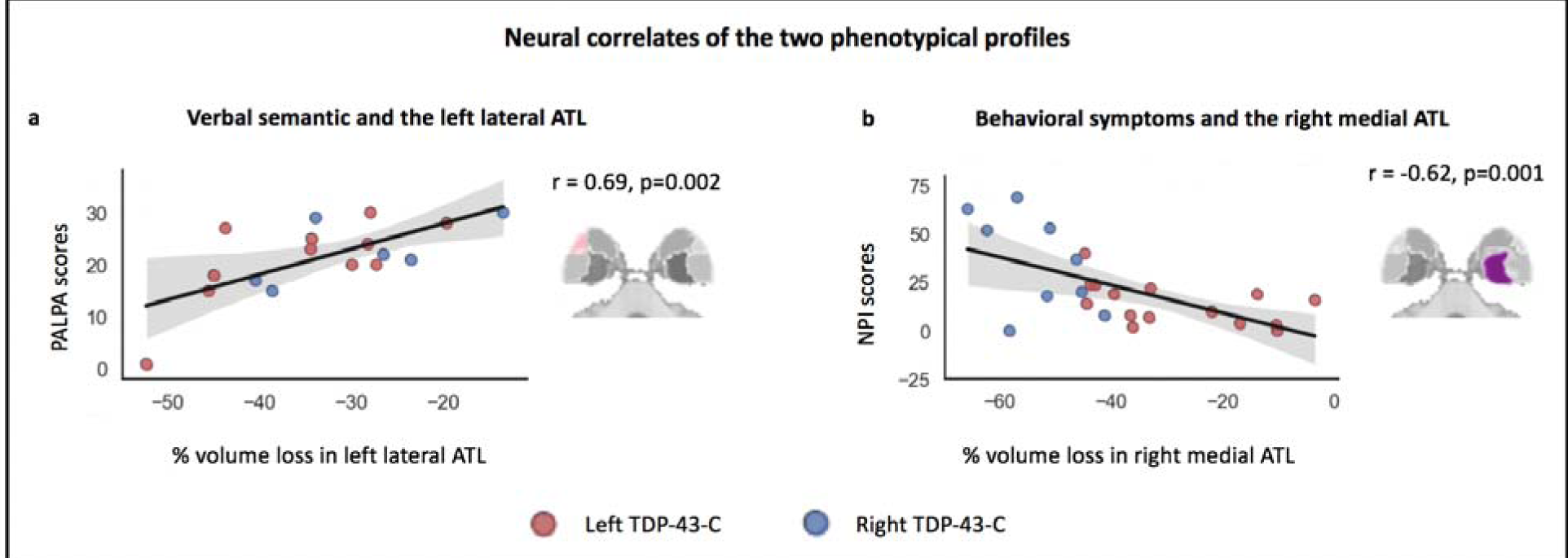
Neural correlates of the two phenotypical profiles. A) Scatter plot illustrating the correlation between verbal semantic deficits (as measured with PALPA irregular words reading, n = 17) and volume loss in the left lateral ATL. B) Correlation between behavioral symptoms scores (as measured with NPI, n = 21) and volume loss in the right medial ATL. Predominantly left cases are depicted with red dots, predominantly right cases with blue ones.

## 4. Discussion

This study leverages the largest cohort of path-proven TDP-43-C cases clinically described, deepening our understanding of regional temporal lobe susceptibility to this proteinopathy and its effects on cognition. We show that predominant right-sided atrophy occurs in up to 40% of cases diagnosed with temporal variants of FTD caused by TDP-43-C. We then provide the first detailed characterization of atrophy distribution within the ATLs, indicating that medial ATL regions are the most vulnerable to TDP-43-C pathology. Finally, we show that atrophy progression is similar irrespective of initial lateralization. Taken together, our findings suggest that, regardless of early clinical and anatomical phenotypical differences, left and right TDP-43-C temporal variants of FTD should be considered spectrum presentation of the same disease.

Hyper-phosphorylated, ubiquitinated and cleaved forms of TDP are considered the histological hallmark of the majority of ubiquitin-positive, tau-, and alpha-synuclein-negative cases of FTLD (known as FTLD- TDP or FTLD-U) (Cairns et al., 2007; Lagier-Tourenne, Polymenidou, & Cleveland, 2010). In the healthy brain, TDP regulates transcription, alternative splicing, binding and stability of RNA (Cohen, Lee, & Trojanowski, 2011; Tollervey et al., 2011), while its pathological depositions are found in neurodegenerative diseases such as FTD and both familial and sporadic amyotrophic lateral sclerosis (ALS) (Neumann et al., 2006; Sreedharan et al., 2008). Different types of TDP pathology have been identified and are currently classified based on the kind of inclusions and their distribution (Mackenzie et al., 2011). Cases with abundance of ubiquitin-positive pathology in upper cortical layers and a preponderance of elongated neuritic profiles over intraneuronal cytoplasmic inclusions are referred to as TDP type C, known in previous classification as type 2 (Mackenzie et al., 2006) or type 1 (Sampathu et al., 2006). Previous reports indicate that in-vivo clinical diagnosis of svPPA is associated with TDP type C in up to 88% of the cases (Josephs et al., 2011; Snowden et al., 2011; Spinelli et al., 2017). Conversely, post-mortem neuropathological diagnosis of TDP-43-C appears to be constantly associated with svPPA presentation (Rohrer et al., 2010; Whitwell et al., 2010). Our study, leveraging the unique sample offered by the UCSF Memory and Aging Center database and brain bank, confirms that post-mortem findings of TDP-43-C pathology are linked with an in-vivo clinical diagnosis of temporal FTD in 95% of the cases (37 out of 39), while a in-vivo clinical diagnosis of temporal FTD is associated with a 84% chance of TDP type C pathology (37 out of 44). More large-scale studies will be needed to elucidate which clinical and/or anatomical findings can help early detection of temporal FTD cases that are due to pathologies other than TDP (16%). Crucially, we add to previous studies by providing strong evidence that early ATL atrophy (irrespective of lateralization) and associated symptoms (verbal and non-verbal semantic and emotional deficits) are the hallmark presentation of TDP-43-C pathology.

Right predominant temporal atrophy was once considered a rare anatomical form of FTD. To our knowledge our study is the first to investigate the prevalence of asymmetric right-sided ATL degeneration in a cohort of homogeneous, path-confirmed FTLD-TDP-C patients. Our findings highlight that right-sided presentations are not uncommon, representing 40% of the total sample, and should always be considered in the differential diagnosis of FTD-spectrum disorders. Nevertheless, our results still show a higher prevalence of left-sided cases (60%). This greater prevalence might not be related to disease vulnerability but rather be explained by a referral bias. It is indeed likely that patients presenting with early behavioral manifestations of right ATL damage would be referred to psychiatric clinics, while left-hemispheric cases with progressive anomia would be promptly evaluated in memory clinics. Consistently, predominantly right cases might reach the attention of neurologists later in the disease course, explaining the less asymmetric presentation of atrophy in these cases. Future studies, integrating psychiatric and neurological knowledge and services, might show that predominantly right and left incipient presentations are equally prevalent. Historically, the first cases of temporal atrophy described were bilateral with slight left-sided asymmetry (Andersen et al., 1997; Hodges, Patterson, Oxbury, & Funnell, 1992). These early cases presented with atrophy already spreading to the posterior ATL and orbitofrontal cortex and thus manifested severe language impairments as well as behavioral changes. Increased awareness of the disorder enabled earlier diagnosis and led to the description of predominantly left vs. right cases (Edwards-Lee et al., 1997; Miller, Chang, Mena, Boone, & Lesser, 1993; Thompson et al., 2003).

The most consistent difference in the clinical presentations is that of more severe anomia in predominantly left cases, detectable even when groups are matched for cross-modal measures of semantic knowledge (Snowden et al., 2017; Woollams & Patterson, 2017). Early reports, classifying patients based on radiologists’ ratings of whole brain CT/MRI scans, suggested a percentage of right-predominant cases around 20% (Snowden, Thompson, & Neary, 2004; Thompson et al., 2003). Subsequent studies systematically quantified the asymmetry of atrophy and suggested that up to 33% of cases might initially present with right-predominant atrophy (Binney et al., 2016). None of the previous work has focused on path-proven cases, however recent advanced subregional segmentation of medial temporal lobe such as the hippocampus and the amygdala has brought attention to an early involvement of the right hemisphere in svPPA (Bocchetta et al. 2019). Using bilateral ROIs, we clustered our sample into left-predominant or right-predominant cases, based on data-driven quantitative analyses of GM volume in the two ATLs. Our results, focusing on the most anterior portion of the temporal lobe, suggest that up to 40% of the cases present with initial right-predominant atrophy, stressing the importance of careful consideration of right ATL during FTD-spectrum disorders differential diagnoses. Coherently, our brain-behavior correlations illustrated by our ROIs, corroborate previous findings establishing the crucial role played by the right temporal lobe in empathic behavior (Perry et al., 2001; Rankin et al., 2006), in particular affect sharing (Shdo et al., 2018), as well as emotions comprehension, especially for negative ones (Rosen, 2002; Rosen et al., 2006).

The biological reasons why the ATLs are vulnerable to FTLD-TDP-C pathology is still unknown. Similarly, the biological bases of different patterns of hemispheric lateralization in different individuals is not known. Interestingly, TDP type C has been associated with a predilection for left temporal structures in primary age-related tauopathy (Josephs et al., 2019), while bilateral temporal involvement is observed in AD (Josephs, 2014), thus further insights might come from a comparison of TDP type C distribution across dementias. Our novel ATL parcellation methodology shows that, irrespective of the hemisphere predominantly affected, atrophy distribution within ATLs describes a gradient whereby medial regions are more affected than lateral ones. These results are consistent with evidence showing that TDP type C pathology spreads from allocortex to neocortex (Nag et al., 2018), a finding supported by in vitro evidence of higher proteotoxicity in the allocortex than in the neocortex (Posimo, Titler, Choi, Unnithan, & Leak, 2013). Animal models have also revealed differences between medial and lateral regions of the temporal pole, with the medial portion presenting the strongest connections to and from the limbic system (Höistad & Barbas, 2008). Recent neuroimaging evidence shows a clear distinction between the pattern of connectivity of medial vs. lateral temporal regions (Bajada, Trujillo-Barreto, Parker, Cloutman, & Lambon-Ralph, 2019). Bajada and colleagues, comparing the connectivity similarities of regions within the temporal lobe, showed that the lateral temporal lobe is characterized by gradual transitions between regions (making it an ideal convergence zone), while the connectivity profile of the medial temporal lobe (in particular that of the hippocampus) is quite heterogeneous. Our observations complement previous reports indicating the same inferior-to-superior gradient, regardless of atrophy lateralization (Binney et al., 2016), strengthening the conclusion that within the ATL pattern of neurodegeneration does not depend on the hemisphere being most affected at initial presentation. Further studies are necessary to investigate specific associations between right or left medial vs. lateral atrophy and different clinical profiles, potentially elucidating the cognitive function subserved by each ATL sub-region. For instance, Vonk and colleagues have described one case in which relative sparing of left dorsolateral ATL might have been sufficient to preserve verbal semantic abilities in a svPPA patient presenting with bilateral but right-predominant atrophy in the medial part of the ATL (Vonk et al., 2019). Taken together, our findings support the hypothesis that TDP-43-C driven neurodegeneration starts in the deep medial temporal structures. Tissue-based studies aiming at understanding the pathophysiology of this disorder should therefore concentrate on these medial regions, regardless of hemispheric lateralization in early disease. Overall, regional susceptibility likely results from the interplay of numerous variables including cell and genetic vulnerability, structural and functional connections to large-scale networks, as well as environmental factors (Miller et al., 2015; Zheng et al., 2018).

Finally, our comparison of the longitudinal evolution of left-predominant and right-predominant cases indicates that atrophy spreads first to the contralateral hemisphere, then to posterior temporal regions and orbitofrontal ones. These findings not only corroborate prior reports in svPPA (Brambati et al., 2009; Kumfor et al., 2016), but also match previously described stages of TDP pathology spread (Brettschneider et al., 2014; Nag et al., 2018; Bocchetta et al., 2020). It has been shown that, across TDP types, pathology first appears localized to the amygdala, then extends to the hippocampus and entorhinal cortex. Subsequently, it spreads to the ATL, eventually involving mid-temporal and orbitofrontal cortices (Nag et al., 2018). A symmetrical spread from orbitofrontal lobes and basolateral amygdala to frontal and temporal lobes, then parietal and occipital lobes has been observed in bvFTD patients (Brettschneider et al., 2014). Only recently, a study focusing on predominantly left TDP-43-C cases, has shown how pathology starts in the left amygdala, left temporal and anteromedial right temporal lobe, then progressively spreads to more posterior and superior parts of the left temporal lobe as well as more lateral regions of the right temporal lobe, finally involving right subcortical structures, and more anterosuperior and basal frontal regions in both hemispheres (Bocchetta et al., 2020). The overall medial-to-lateral, frontal-to-posterior trend of neurodegeneration caused by TDP-43 seems thus to hold for TDP-43-C cases, yet with the key characteristic of starting in medio-temporal (rather than medio-frontal) structures. This finding, which needs to be corroborated by studies comparing predominantly left and right TDP type C cases, could prove critical for pathological early differential diagnosis. Finally, these neuroanatomical observations have important clinical implications, as they are parallel by neuropsychological findings indicating a progressive overlap of the clinical syndromes, being virtually indistinguishable within 3 years from diagnosis (Kumfor et al., 2016; Seeley et al., 2005).

Our findings of bilateral temporal lobe susceptibility to TDP-43-C has important implications for clinical practice in terms of both diagnosis and treatment. Currently, most patients are diagnosed as svPPA if they meet consensus criteria for PPA (Mesulam, 2003), and specific criteria for the semantic variant (Gorno-tempini et al., 2011). The latter must show impaired confrontation naming and impaired single-word comprehension, together with at least three of the following: impaired object or face knowledge, surface dyslexia or dysgraphia, spared repetition, or spared speech production. As suggested by previous reports and supported by our findings, left-predominant cases usually meet these criteria, while the diagnosis of right-predominant cases is more difficult, because there are no specific criteria for the right variant of temporal FTD. These patients might be diagnosed with bvFTD if the first (and predominant) symptoms include at least three of the following: behavioral disinhibition, apathy or inertia, loss of sympathy or empathy, perseverative or compulsive behavior, hyperorality and dietary changes, or executive deficits (Rascovsky et al., 2011). As described earlier, this is often the case. Alternatively, they might be assigned the label of the right variant svPPA if nonverbal semantic deficits are investigated and detected, even though consensus criteria for such a variant do not currently exist. It should be noted that, as in our sample, most of these right-predominant patients would meet Neary criteria for SD (1988) and svPPA (anomia and semantic deficits for objects or faces) but not Mesulam criteria for general PPA, as language deficits might not be the first complaint. Finally, many patients with right-predominant temporal variant of FTD were first referred to psychiatric care when behavioral symptoms are preponderant and not framed in the context of a progressive deterioration of the cognitive profile (Mendez & Perryman, 2002). Certainly, the clinical profile of svPPA and bvFTD patients greatly overlaps (Blair, Marczinski, Davis-Faroque, & Kertesz, 2007). Language impairments are not rare in bvFTD patients (Hardy et al., 2016), and only subtle behavioral symptoms analyses can discriminate right temporal from right frontal FTD cases: the former show increased mental rigidity and depression, while the latter exhibit greater disinhibition (Bozeat, Gregory, Lambon Ralph, & Hodges, 2000). Overall, the diagnosis of bvFTD appears to be one of the most difficult and least stable over time within the FTD spectrum (Perry et al., 2019). Separating the right-temporal variant from bvFTD might help simplify this complex scenario.

Overall, our results suggest that TDP-43-C driven temporal variant FTD should be suspected when patients’ clinical presentation includes either symptoms indicative of a left ATL involvement, such as poor confrontation naming and single word comprehension, or indicative of right ATL damage, such as a lack of socioemotional sensitivity or empathy and loss of semantic knowledge for faces and known people. If neuro-anatomical data are available, clinicians should assess the degree of asymmetric ATL involvement, and whether the gradient is medial-to-lateral. This can inform on which additional cognitive and behavioral manifestations are to be expected. It should be stressed that while svPPA/temporal variant FTD are overwhelmingly associated with TDP-43-C, bvFTD presents considerably more pathological variability (Perry et al., 2017). The identification of in vivo features that predict a pathological diagnosis is increasingly important, as pharmacological trials targeting specific proteinopathy emerge. In the future, diagnostic help could come from [18F]AV-1451 PET (so called TAU PET) which seems to bind to bilateral temporal lobe pathology in svPPA cases, but does not show frontal/temporal binding in bvFTD (Bevan-Jones et al., 2018; Josephs et al., 2018; Makaretz et al., 2018). However, to date no biomarker for TDP exists (Steinacker, Barschke, & Otto, 2018), so the integration of clinical and neuroimaging findings is still the best way to diagnose these patients. Regarding therapy trials, the implications of our findings are two-folds. First, information on the most likely underlying proteinopathy is crucial to decide which patients to include, and our results suggest that all patients presenting with temporal variant of FTD, irrespective of atrophy lateralization, should be treated as highly probable TDP-43-C cases (and thus included for drugs targeting TDP). Second, knowledge of the longitudinal atrophy progression patterns suggests which cortical regions should be monitored to assess whether a particular treatment is slowing the spread of atrophy. Atrophy progression should be tracked by mapping the progressive involvement of the ipsilateral lateral ATL, ipsilateral posterior temporal areas, and contralateral hemisphere.

Despite the robust size of this pathology-proven cohort, future studies including more subjects are warranted. Better powered studies will be instrumental in establishing whether current investigation failed to detect interactions between disease epicenters (i.e., left vs. right anterior temporal pole) and atrophy distribution (i.e., from medial to lateral). Moreover, this retrospective, path confirmed study includes relatively old cases, so the neuropsychological data available do not include more recent measures such as Famous Faces Naming (Borghesani et al., 2019) or emotion processing tasks (Kumfor et al., 2018). Larger samples, including the more targeted neuropsychological tests now available, will allow more refined investigation of the neural correlates of specific phenotypical characteristics of TDP-43-C driven temporal FTDs. One important caveat of the present study is the lack of an in-depth analysis of the two MCI cases with post-mortem findings of TDP-43-C pathology. Future investigations of the anatomical and neuropsychological characteristics of such cases will speak to the generalizability of our observations and be instrumental in elucidating the features of prodromal left- and right-sided temporal FTD. Finally, building on our findings, future studies should aim at comparing predominantly right temporal FTD cases with bvFTD ones. This comparison would benefit from careful assessment of social, emotional, and nonverbal semantic processing as well as from the inclusion, as ROIs, of extra-temporal regions (e.g., the orbitofrontal cortex) and subcortical ones (e.g., amygdala and insula). Ultimately this kind of studies will lead to updated clinical criteria for temporal variants of FTD able to capture both language and behavioral symptoms.

In conclusion, we showed that TDP-43-C associated FTD cases might present with predominantly right ATL atrophy in up to one third of cases, yet exhibit the same atrophy distribution within, and longitudinal spread outside, the ATLs. Specifically, the medial portion of the ATLs appears to be particularly susceptible to TDP-43-C driven neurodegeneration, irrespective of incipient atrophy lateralization. Hence, while the different linguistic and behavioral features of the two presentations will require different symptomatic and supportive strategies, both left and right phenotypes of temporal FTD should be considered the same disorder from a molecular perspective.

## Acknowledgements

The authors thank the patients and their families for the time and effort they dedicated to the research. We thank Anna Karydas and Jennifer Yokoyama (UCSF Memory and Aging Center genetics experts) for assistance with genetic data, as well as Dr. John Trojanowski (Department of Pathology and Laboratory Medicine, University of Pennsylvania, Philadelphia) for assistance with autopsy data. The authors declare no competing financial interests.

